# ARF small GTPases in the developmental function mediated by ARF regulators GNOM and VAN3

**DOI:** 10.1101/2022.01.07.475425

**Authors:** Maciek Adamowski, Ivana Matijević, Jiří Friml

## Abstract

ARF small GTPases are molecular switches acting in intracellular trafficking. Their cycles of activity are controlled by regulators, ARF Guanine nucleotide Exchange Factors (ARF-GEFs) and ARF GTPase Activating Proteins (ARF-GAPs). The ARF-GEF GNOM (GN) and the ARF-GAP VAN3 share a prominent function in auxin-mediated developmental patterning, but the ARFs which they might control were not identified. We conducted a loss-of-function and localization-based screening of the *ARF/ARF-LIKE* gene family in *Arabidopsis thaliana* with the primary aim of identifying functional partners of GN and VAN3, while extending the limited understanding of this gene group as a whole. We identified a function of ARLA1 in branching angle control. Mutants lacking the variably localized ARLB1, ARFB1, ARFC1, ARFD1, and ARF3, even in high order combinations, do not exhibit any evident phenotypes. Loss of function *arfa1* phenotypes support a major role of ARFA1 in growth and development overall, but patterning defects typical to *gn* loss of function are not found. ARFA1 are not localized at the plasma membrane, where GN and VAN3 carry out developmental patterning function according to current models. Taken together, putative ARF partners of GN and VAN3 in developmental patterning cannot be conclusively identified.

## Introduction

ARF small GTPases function in eukaryotic endomembrane systems as molecular switches recruiting various effector proteins to membranes in a highly controlled manner (reviewed in Donaldson and Jackson, 2011; Jackson and Bouvet, 2014; Singh and Jürgens, 2017; Yorimitsu et al., 2014). The reversible activity of ARFs depends on their switching between GDP and GTP-bound forms. ARF-GDP preferentially remain cytosolic and typically inactive, while ARF-GTP are membrane-docked and active. Active ARF-GTP recruit effectors, among which prominent are components of vesicle coats (Paczkowski et al, 2015). A plethora of effectors, including many non-coat components, is known in non-plant systems (Khan and Ménétrey, 2013; Cherfils, 2014). ARF cycles are controlled by two classes of regulatory proteins. The hydrolysis of GTP by the ARF GTPase is stimulated by ARF-GAPs (ARF GTPase Activating Proteins), while the replacement of GDP with GTP is promoted by ARF-GEFs (ARF Guanine nucleotide Exchange Factors).

In *Arabidopsis thaliana*, the study of the ARF machinery has been primarily focused on the ARF regulators. A number of characterized GEFs and GAPs perform the conserved, fundamental trafficking functions. The ARF-GEF GNOM-LIKE1 (GNL1) of the GBF1 class has a function in secretion at the Golgi apparatus (GA) which it shares with its homologue GNOM (GN) (Richter et al., 2007; Teh and Moore, 2007). ARF-GEFs of the BIG class localized at the *trans-*Golgi Network (TGN) are required for exocytosis and trafficking to the cell plate (Richter et al., 2014). Among ARF-GAPs, named AGD (ARF GAP DOMAIN) in *A. thaliana*, AGD6,7 and AGD8-10 subfamilies, homologous to non-plant ARFGAP1/Gcs1p and ARFGAP2,3/Glo3p, respectively, act non-redundantly in COPI-coated vesicle formation at the GA (Min et a., 2007, 2013), while AGD5/NEVERSHED/MTV4 (MODIFIED TRANSPORT TO THE VACUOLE4) is involved with clathrin-coated vesicles formed at the TGN (Stefano et al., 2010; Sauer et al., 2013).

Prominently, some of the ARF regulators in plants are involved in developmental body patterning connected with the function of the hormone auxin. The activity of the ARF-GEF GN is linked with the control of the polar distribution and transport activity of PIN auxin efflux carriers (Steinmann et al., 1999; Geldner et al., 2003; Adamowski and Friml, 2015). Loss of function manifests in embryonic and post-embryonic patterning defects (Mayer et al., 1991, 1993; Shevell et al., 1994; Geldner et al., 2004; Okumura et al., 2013; Verna et al., 2019). Recent findings indicate that the developmental function of GN is mediated from structures of unknown nature localized at the plasma membrane (PM) (Adamowski et al., 2022). The ARF-GAP VAN3 (VASCULAR NETWORK DEFECTIVE3), also known as SFC/FKD2/AGD3 (SCARFACE/FORKED2), and its homologues, function in auxin-mediated development similarly to GN, where their role is especially visible in the formation of vascular networks in cotyledons and leaves (Koizumi et al., 2005; Sieburth et al., 2006). VAN3 too, acts from the PM, but its localization pattern there differs from GN (Naramoto and Kyozuka, 2018; Adamowski and Friml, 2021).

In contrast to these GEF and GAP regulators, the ARF small GTPases of *A. thaliana* have been studied in less detail. Conclusive evidence for the participation of specific ARF/ARLs in the developmental patterning function characteristic for GN and VAN3 is lacking. The *ARF/ARF-LIKE (ARL)* gene family of *A. thaliana* contains 19 genes (Singh et al., 2018). Several loss-of-function mutants were isolated, notably the embryo defective *arlc1/hallimasch/titan5* (Mayer et al., 1999; McElver et al., 2000), while other *ARF* members were studied by post-translational silencing (Gebbie et al., 2005). In other cases, dominant mutant variants that are preferentially GDP- or GTP-bound were employed to deduce function (e.g. Xu and Scheres, 2005). Some of the ARF proteins were localized to the GA, TGN, and PM, by immunoelectron microscopy and fluorescent protein fusion approaches, sometimes with the use of transient expression systems in protoplasts (e.g. Robinson et al., 2011, Matheson et al., 2008; Singh et al., 2018). Indirect lines of evidence indicate that the *ARFA1* group may be responsible for all essential membrane traffic (Singh et al., 2018), but overall, relatively little is understood about functions of the large family of *A. thaliana* ARF/ARL small GTPases.

Here, we report on a study of the *ARF/ARL* gene family in *A. thaliana*, focused in particular on genetic knock-outs and *in planta* fluorescent protein fusion localization approaches. Our primary goal was to identify ARF/ARL partners that may engage with the ARF regulators GN and VAN3 as mediators of their developmental patterning function, but the study provides insights and genetic materials relevant to all aspects of *ARF/ARF-LIKE* gene function.

## Results

### Loss-of-function and localization-based screening of the *ARF/ARL* gene family

The *ARF/ARL* gene family of *A. thaliana* consists of 19 genes (Singh et al., 2018). We undertook to characterize this gene family through an extensive loss-of-function study, aided by subcellular localization data, by gene expression data from published databases (Winter et al., 2007; Klepikova et al., 2016), and by a summary of previous research. We characterized the family members with a primary goal of identifying ARF/ARL proteins that may function with GN and VAN3 in the developmental patterning role of these ARF regulators. Our reasoning is that potential ARF/ARL candidates are likely to be found at the PM, from where GN and VAN3 act based on recent models (Naramoto and Kyozuka, 2018; Adamowski et al., 2022), and that the corresponding mutants are likely to exhibit developmental phenotypes similar especially to *gn*, where GEF-mediated activation of the presumed target ARFs is missing. Given that ARLs, named so due to a limited homology to prototypical ARFs, can in some instances be activated by the action of GEFs, like ARFs (Chen et al., 2010), and given that some small GTPase isoforms of *A. thaliana* were variably designated as ARF or ARLs (e.g. ARFD1; Nielsen et al., 2006; Singh et al., 2018), we studied the whole *ARF/ARL* gene family of *A. thaliana* indiscriminately. Suppl. Table 1 contains a list of *ARF/ARL* genes in *A. thaliana* and indicates the nomenclature used in this study, as well as aliases used elsewhere, and contains details of the generated loss-of-function alleles in the individual genes. The isolated mutants are strictly knock-out alleles, harbouring T-DNA insertions, or CRISPR/Cas9-generated insertions and deletions, in exons of the *ARF/ARL* genes.

### Characterization of *ARLA1, ARLB1*, and *ARLC1*

The *ARLA1* gene group consists of four members, *ARLA1a-ARLA1d*. The function of these genes is poorly understood, but ARLA1 proteins are employed by *Tobamovirus* for the replication of its genomic RNA both in *Nicotiana tabacum* and in *A. thaliana* (Nishikiori et al., 2001). The genes have distinct expression patterns (Winter et al., 2007; Klepikova et al., 2016): *ARLA1a* and *ARLA1d* are ubiquitously expressed but reach peak expression in seeds and mature pollen, respectively, while *ARLA1b* is relatively specific to the seedling root, and *ARLA1c* to the anthers of mature flowers. *ARLA1b* may be a pseudogene, as the predicted protein lacks an N-terminal part conserved in other ARLA1 proteins (Nishikiori et al., 2001).

We generated stably expressed GFP fusions of the ARLA1 sequences driven by the *UBQ10* promoter *(UBQ10*_*pro*_*:ARLA1a-GFP* to *UBQ10*_*pro*_*:ARLA1d-GFP)* and using confocal laser scanning microscopy (CLSM), observed that all ARLA1 proteins localize to the cytosol when expressed in seedling root apical meristem (RAM) epidermis, while no association to membrane structures could be seen (Figure 1A). Using CRISPR/Cas9, we generated an *arla1a arla1b arla1c arla1d (arla1abcd)* mutant. The mutant seedlings developed normally (Figure 1B), while the adult plants were characterized by wide angles of lateral branches, and of siliques (Figure 1C). This phenotype was not present in the *arla1bc* double mutant (Suppl. Figure 1A). The shoot phenotype of *arla1abcd* might indicate a function of *ARLA1* in the gravitropic setpoint angle control (Digby and Firn, 1995; Roychoudhry and Kepinski, 2015) or might be explained by deficiencies in cell wall rigidity. While the finding that the *ARLA1* gene family influences branching angles in the shoot is a valuable insight, both the localizations of the proteins, and the overall mutant phenotype do not support the notion that ARLA1 group engages with GN or VAN3 in their developmental function.

**Figure 1.**
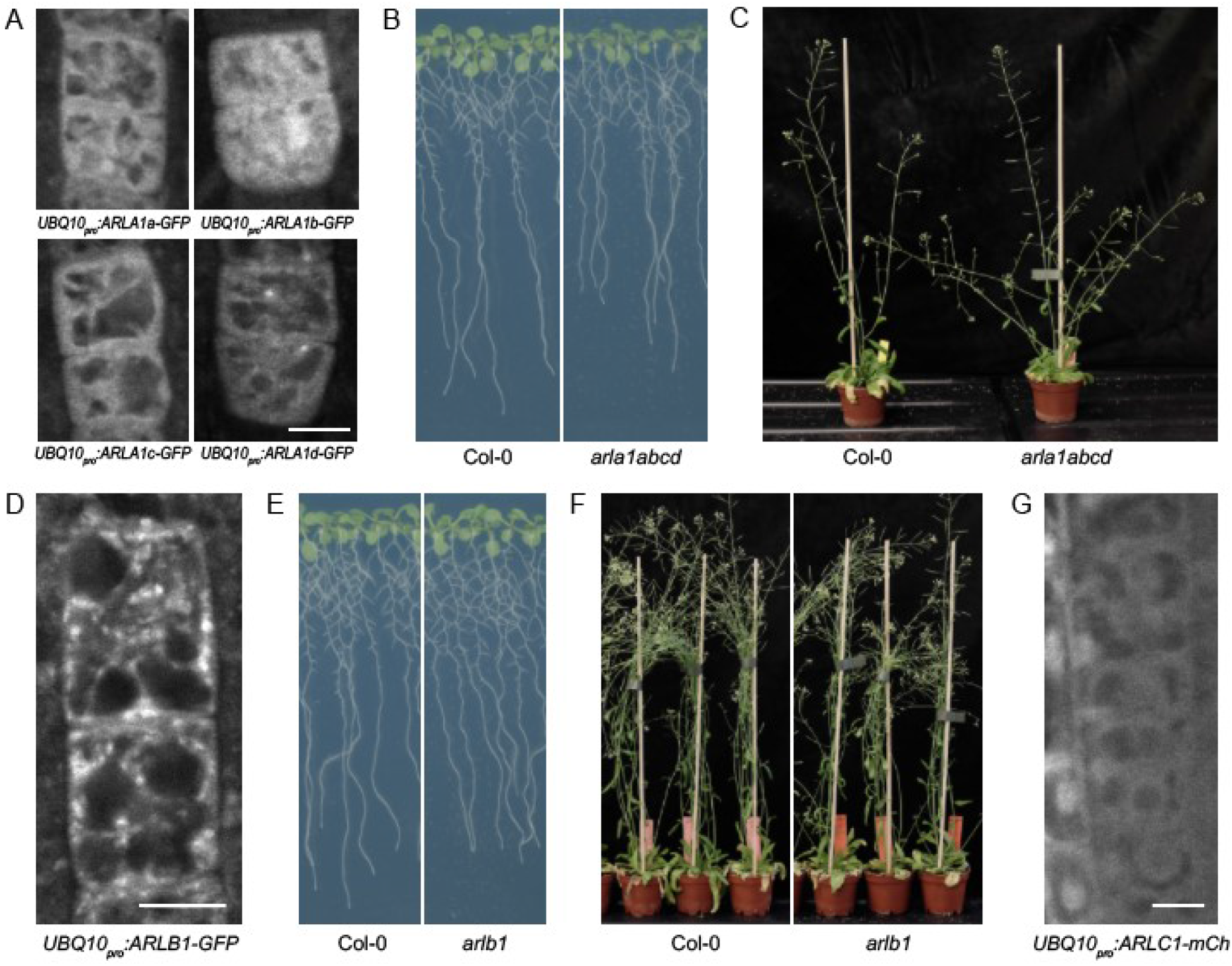
Characterization of ARLA1, ARLB1, and ARLC1. (A) CLSM images of ARLA1-GFP protein fusions in the epidermis of RAMs. ARLA1 proteins localize to the cytosol. Punctate signals in the ARLA1d-GFP panel are most likely autofluorescent structures detected due to low expression of the fluorescent protein fusion. Scale bar - 10 μm. (B) Normal development of arla1bcd mutant seedlings in an in vitro culture. (C) Adult aria1abcd mutants are characterized by wide branching angles of shoots and of siliques. (D) A CLSM image of ARLB1-GFP protein fusion in the RAM epidermis. ARLB1 localizes to endomembrane structures. Scale bar - 10 μm. (E) Normal development of arlb1 mutant seedlings in an in vitro culture. (F) Normal development of arlb1 adults. (G) A CLSM image of ARLC1-mCherry protein fusion in the RAM. ARLC1 localizes to the cytosol and to nuclei.

**Suppl. Figure 1.**
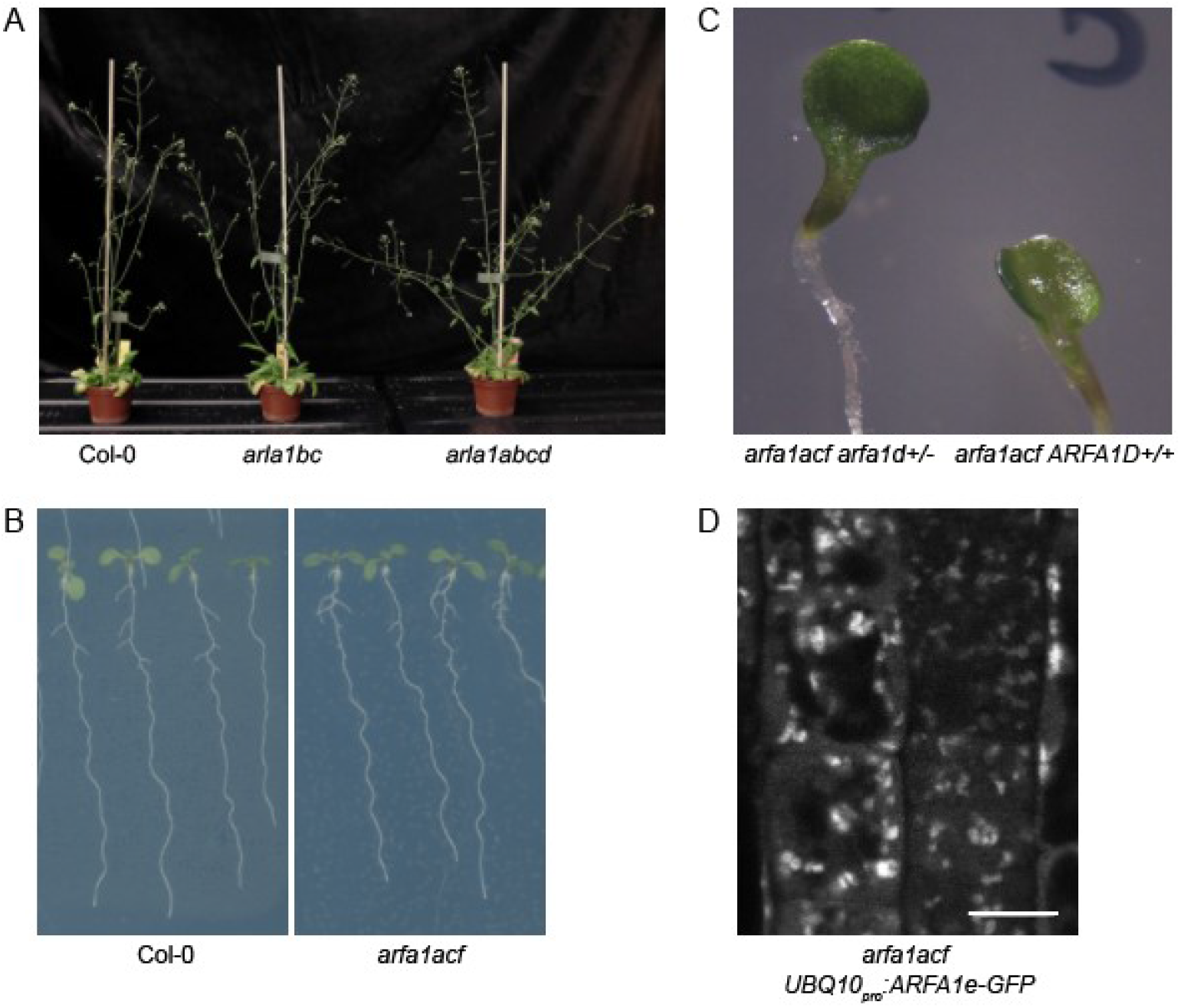
Additional data related to ARF and ARL characterization. (A) Adult arla1bc mutants do not exhibit the branching phenotype of arla1abcd. (B)Seedings of arfa1acf mutants appear normal. (C) Seedings with single cotyledonsfound rarely in a segregating population of arfa1acf d+/-. Genotypes of the seedings are shown. (D) A CLSM image pf ARFA1e-GFP in the seeding RAM of arfa1acf showing localization to aggregated endosomal structures.

*ARLB1* is a ubiquitously expressed single copy gene in *A. thaliana* with an uncharacterized function. We generated a *UBQ10*_*pro*_*:ARLB1-GFP* reporter line and observed that ARLB1 localizes to punctate structures in the endomembrane system, but not to the PM (Figure 1D). An *arlb1* T-DNA mutant developed normally both at seedling and at adult stages (Figure 1E and 1F). At present, no function can be ascribed to the *ARLB1* gene.

*ARLC1* is a single copy gene described on the basis of the isolation of embryo defective mutants *titan5* and *hallimasch* (Mayer et al., 1999; McElver et al., 2000). Mutants are characterized by a failure in cell divisions, but not in cell cycle progression, resulting in embryos and endosperm composed of very large cells with enlarged nuclei (Mayer et al., 1999; McElver et al., 2000). The function of ARLC1 is likely connected with the control of microtubule cytoskeleton (Mayer et al., 1999). The expression of *ARLC1* extends beyond embryonic development (McElver et al., 2000; Winter et al., 2007; Klepikova et al., 2016). When expressed in the seedling RAM, an ARLC1-mCh fusion protein localized to the cytosol and to nuclei (Figure 1G). The embryonic mutant phenotype, which appears to affect processes distinct from those where GN and VAN3 are involved, as well as the localization data, make *ARLC1* an unlikely candidate for the participation in molecular processes controlled by these ARF regulators.

### Characterization of *ARFB1, ARFC1, ARFD1, and ARF3*

Next, we characterize a group of a total of 7 genes named *ARFB1, ARFC1, ARFD1* and *ARF3*, placed on phylogenetic trees between the above-described *ARLA1, ARLB1*, and *ARLC1* genes, and the *ARFA1* group (Singh et al., 2018). Of these, *ARFC1, ARFD1*, and/or *ARF3* are classified as *ARL*, rather than *ARF*, by some authors (Latijnhouwers et al. 2005; Stefano et al., 2006; Nielsen et al., 2006; Singh et al., 2018). By our choice of nomenclature we do not intend to group the genes into one, or other, class.

Limited information suggests distinct characteristics of these genes. The *ARFB1a-ARFB1c* genes were originally grouped into a common class, but subsequent phylogenetic and localization studies suggest their division into two distinct classes (*ARFB1a* and *ARFB1b,c;* Singh et al., 2018). *ARFB1a* has been localized to the PM and TGN (Matheson et al., 2008; Singh et al., 2018), while *ARFB1b* to the GA and TGN (Singh et al., 2018). All three genes are ubiquitously expressed, but are up-regulated in senescent leaves (Winter et al., 2007; Klepikova et al., 2016). An artificial microRNA (amiRNA) line down-regulating *ARFB1a* expression, an *arfb1b arfb1c* double knockout, or the cross of these, do not exhibit evident developmental phenotypes (Singh et al., 2018). *ARFC1* is encoded by a single gene with a high and ubiquitous expression. Recently, a mutant in this gene was identified in the *modified transport to the vacuole (mtv)* forward genetic screen as defective in vacuolar trafficking (Delgadillo et al., 2020). The ARFC1 protein was localized to the *trans* side of GA in a transient expression system (Delgadillo et al., 2020). *ARFD1* is an ARF clade present only in crucifers (*Brassicaceae;* Singh et al., 2018). In *A. thaliana* it comprises two genes, *ARFD1a* and *ARFD1b*, whose expression is preferential to the seedling root (Winter et al., 2007; Klepikova et al., 2016). Finally, *ARF3*, most likely a homologue of the yeast *Arl1* (Yu and Lee, 2017), is a ubiquitously and highly expressed (Winter et al., 2007; Klepikova et al., 2016) single copy gene, whose product has been localized to the GA, where it functions in the recruitment of the golgin AtGRIP, a protein involved in the maintenance of GA structure (Latijnhouwers et al. 2005; Stefano et al., 2006).

To study this ARF/ARL group, we generated a set of stably transformed fluorescent reporter lines controlled by the *UBQ10* promoter, or, when this approach did not lead to expression, we employed the estradiol induction system instead. With CLSM, we confirmed that ARFB1a localizes to the PM, and variably observed it at endosomal structures, as well (Figure 2A). In Total Internal Reflection Fluorescence (TIRF) microscopy, the PM pool of ARFB1a presented a punctate pattern of very dynamic signals (Figure 2B and Suppl. Movie 1). This localization was not reminiscent of the typically stable, and more scarce, PM-localized structures containing GN (Adamowski et al., 2022), but was similar to the dynamic PM associations of VAN3 (Adamowski and Friml, 2021). ARFB1b, ARFB1c, ARFD1a, and ARF3 were observed with CLSM in punctate structures in the cytosol, likely organelles of the endomembrane system (Figure 2A), but their exact nature was not established. ARFC1 was only present in the cytosol (Figure 2A). The difference between this observation and the localization of ARFC1 to the *trans* Golgi side described previously (Delgadillo et al., 2020) may be due to the very low expression in our *UBQ10*_*pro*_*:ARFC1-GFP* lines.

**Figure 2.**
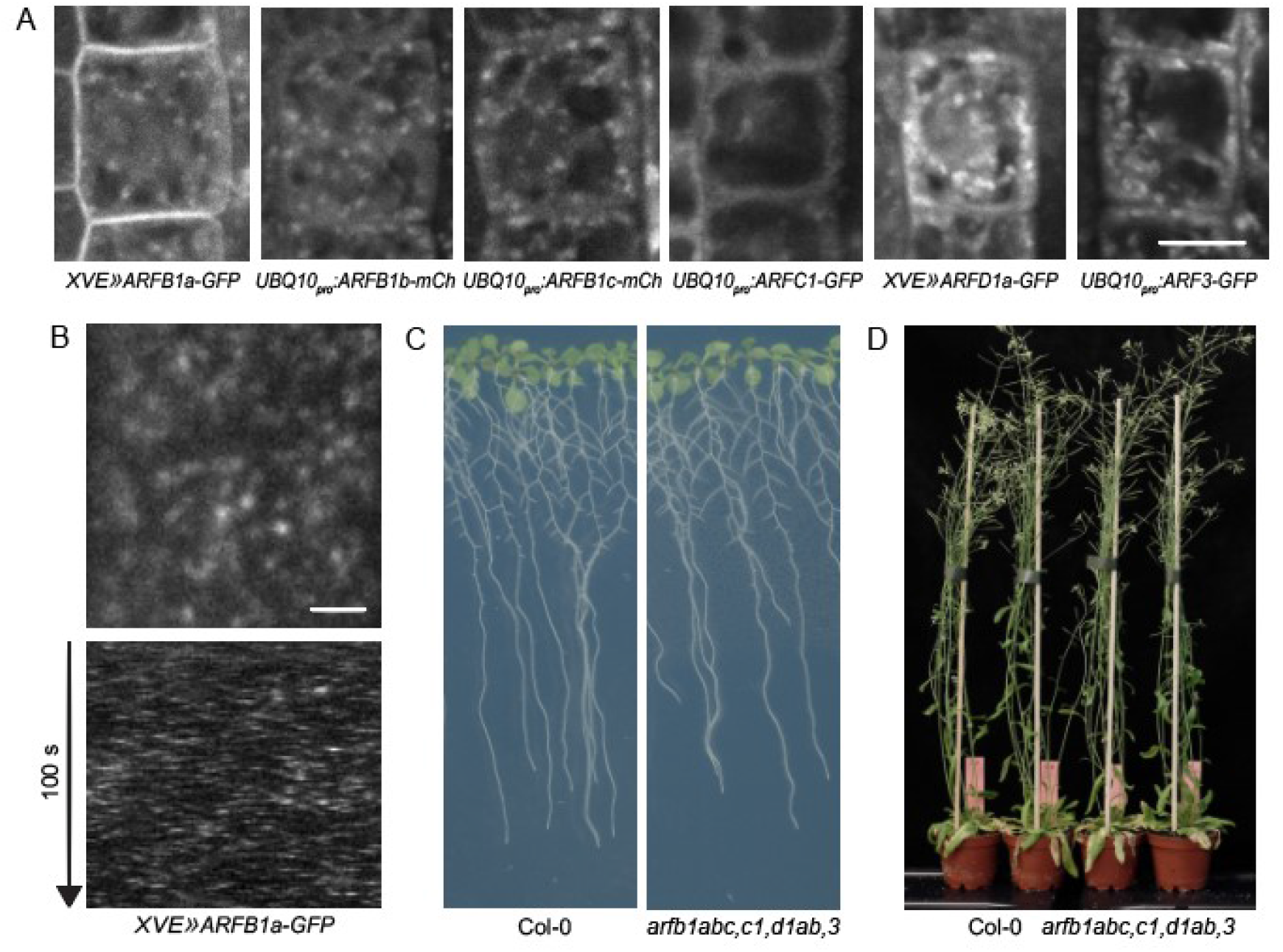
Characterization of ARFB1, ARFC1, ARFD1, and ARF3. (A) CLSM images of ARFB1, ARFC1, ARFD1, and ARF3 fluorescent protein fusions in RAM epidermis. ARFB1a-GFP localizes to the PM and variably to endosomal structures. ARFB1b-mCherry, ARFB1c-mCherry, ARFD1a-GFP and ARF3-GFP localize to endosomal structures. ARFC1-GFP localizes to the cytosol. Scale bar - 10 μm. (B) TIRF Image (upper panel) and kymograph of a TIRF movie (bottom panel) of ARFB1a-GFP in etiolated hypocotyl epidermis. ARFB1 a-GFP localizes to the PM as very dynamic punctate structures. Scale bar - 2 μm. See also Suppl. Movie 1. (C) Normal development of arfb1abc,c1,d1ab,3 mutant seedlings in an in vitro culture. (D) Normal development of arfb1abc,c1,d1ab,3 adults.

To test the function of these genes, we first generated *arfb1abc, arfd1ab*, and *arf3* knockout mutants by CRISPR/Cas9 mutagenesis, and isolated a T-DNA *arfc1* mutant allele since named *arfc1-2* (Delgadillo et al., 2020). None of these mutants exhibited visible morphological phenotypes (not shown). Considering that the genes from these *ARF/ARL* groups are neighbours on phylogenetic trees (Singh et al., 2018), and that even functionally distinct ARF and ARL proteins can possess some shared effectors (Van Valkenburgh et al., 2001), we opted to generate a complete mutant of this gene group, to evaluate a potential functional redundancy. The resulting *arfb1abc arfc1-2 arfd1ab arf3* knockout mutant was also phenotypically normal, during both seedling and adult stages of development (Figure 2C and 2D).

In summary, we do not find significant evidence that *ARF/ARL* genes from the *ARFB1, ARFC1, ARFD1* and *ARF3* classes are involved in the function associated with the ARF regulators GN and VAN3. The localization of ARFB1a to the PM, where it dynamically associates in a manner similar to VAN3, is the only indication of a possible functional link. Yet, loss of function analysis, including high order mutants, strongly argues against the involvement of *ARFB1a* or other genes from this group in developmental patterning. The functions of these genes, which in any case appear quantitatively not major, are at present not elucidated.

### *arfa1* loss-of-function phenotypes

Mutant phenotypes and localization patterns do not support any of the ARF/ARLs analyzed so far as potential functional partners of GN and VAN3. At this stage, the only group remaining are *ARFA1*. The *ARFA1* group consists of 6 members, *ARFA1a-ARFA1f*, encoding proteins of very high sequence similarity. Two of the genes, *ARFA1a* and *ARFA1d*, encode identical proteins. With the exception of the pollen-specific isoform *ARFA1b*, the *ARFA1* genes are expressed ubiquitously (Winter et al., 2007; Klepikova et al., 2016). Expression of mutant ARFA1 variants deficient in GDP to GTP exchange, or in GTP hydrolysis, interferes with several trafficking pathways and affects endomembrane system integrity (Lee et al., 2002; Takeuchi et al., 2002; Pimpl et al., 2003; Xu and Scheres, 2005; Singh et al., 2018). The developmental function of this subfamily has been studied by an RNA interference (RNAi) approach, where down-regulation of *ARFA1* expression lead to stunted growth, reduced cell production rates and final cell sizes, alteration of flowering time and apical dominance, and reduced fertility (Gebbie et al., 2005). Another study using dominant variants described a function of *ARFA1* in epidermal cell polarity, manifested by the positioning of root hair outgrowths (Xu and Scheres, 2005). A dominant variant of *ARFA1c* was found in a forward genetic screen related to trafficking of PIN1 auxin efflux carriers (Tanaka et al., 2014). Singh et al. (2018) propose that the *ARFA1* class is responsible for all significant ARF-related trafficking function, including the activity of GN. ARFA1 localization is well defined by immunoelectron microscopy studies with antibodies raised against ARFA1c: These small GTPases localize to the GA and TGN (Robinson et al., 2011).

To assess a potential function of the ARFA1 group as mediators of the developmental function of GN, we generated *arfa1* mutants using CRISPR/Cas9. In comparison to the previous efforts based on inducible post-transcriptional silencing (Gebbie et al., 2005), we hoped to observe the consequences of *arfa1* loss of function in embryonic development, where GN prominently acts, and to potentially obtain a higher degree of *arfa1* deficiency. With CRISPR/Cas9, we targeted all *ARFA1* genes except the pollen specific isoform *ARFA1b*. Initially, we isolated an *arfa1e* single mutant, which was phenotypically normal (Figure 3A), and an *arfa1acf* triple mutant, which presented a mild decrease in overall growth as an adult, but developed normally at the seedling stage (Figure 3A and Suppl. Figure 1B). In a second round of mutagenesis, we targeted *ARFA1d* and *ARFA1e* in the *arfa1acf* background. The *arfa1acfe* homozygote was characterized by a stunted growth of adults and a very limited fertility (Figure 3A). In contrast, *arfa1acfd* mutants could be maintained only in a segregating state, as the homozygotes did not develop into maturity. Quintuple *arfa1acfde* mutants could not be isolated: *arfa1acfe* seedlings carrying the CRISPR/Cas9 transgene able of producing mutations in *ARFA1d* never had an *arfa1d* mutation, while individuals of the genotype *arfa1acf* -/-*arfa1d* +/-*arfa1e* +/-were invariably infertile and their progenies could not be analyzed. Gene dosage effects associated with the *arfa1d* mutation in a heterozygous state were also seen in a slight, but observable, phenotypic difference between adult individuals of genotypes *arfa1acf* -/- and *arfa1acf* -/- *arfa1d* +/- (not shown).

**Figure 3.**
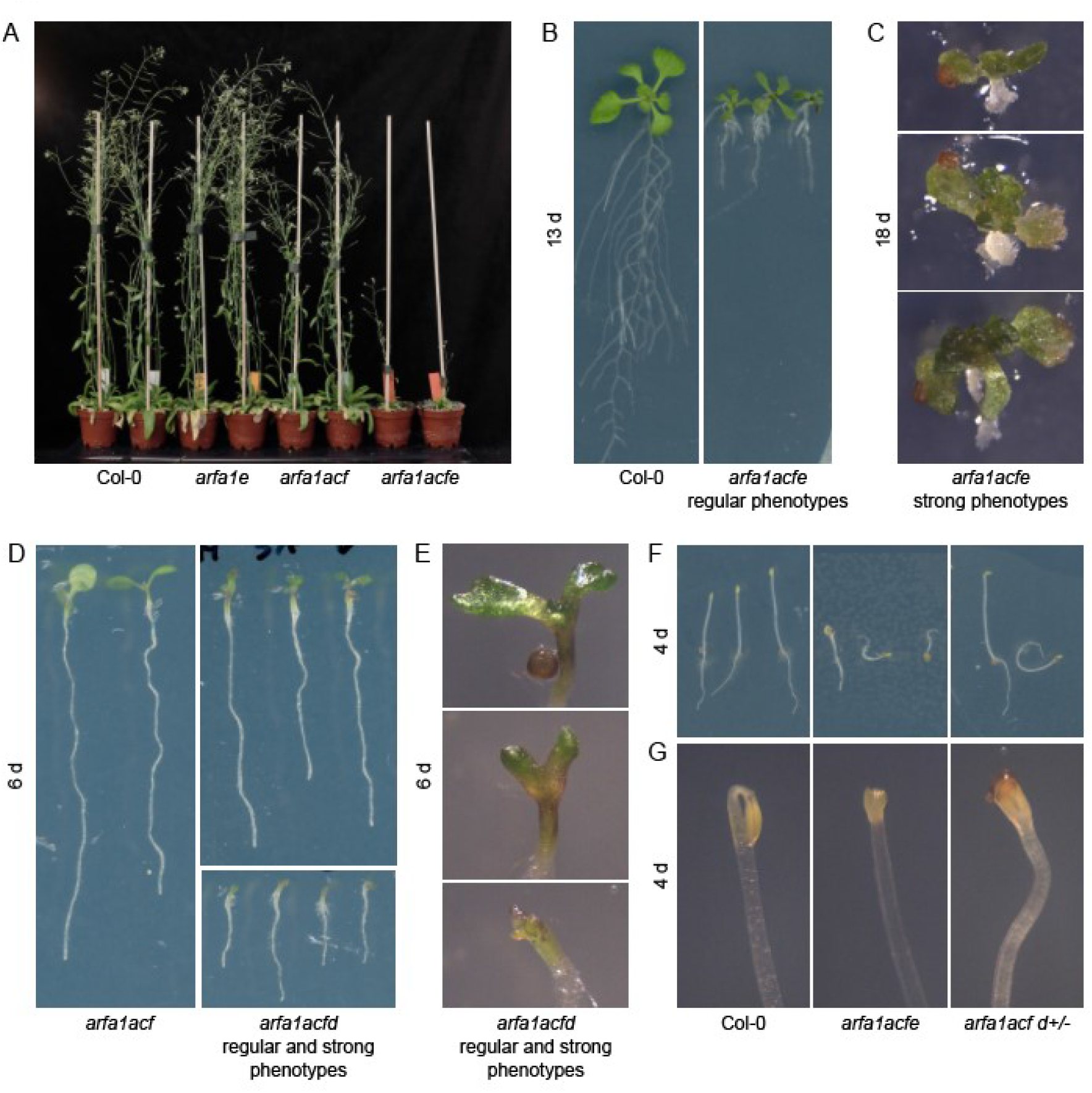
arfa1 loss-of-function phenotypes. (A) Adult arfa1 mutants, arfa1e are phenotypically normal, arfa1acf exhibit a mild reduction in growth, and arfa1acfe are have strongly reduced growth and low fertility. (B) Seedling phenotypes of arfa1acfe mutants of regular phenotypic strength. (C) Strong seedling phenotypes of arfa1acfe mutants characterized by an arrest of root development but no defects in apical patterning (D) Regular and strong phenotypes of arfa1acfd mutants (E) Detail of apical development of regular and strong arfa1acfd mutants. The mutation leads to apical deletion in strong cases, while weaker phenotypes are characterized by development of two cotyledons. (F) Etiolated seedlings of arfa1acfe and arfaf1acf d+/- lines. Seedlings growing agravitropically are observed in both mutant lines. (G) Apical hooks of arfa1acfe and arfa1acfd are often open after 4 d of development.

Homozygous *arfa1acfe* seedlings exhibited a range of phenotypes. Seedlings with regular phenotypes, some of which could develop into adults, had severely retarded growth in *in vitro* cultures (Figure 3B). *arfa1acfe* mutants of strongest phenotypic classes had arrested roots and strongly misshapen cotyledons and leaves (Figure 3C). Yet, even these severely defective seedlings were a product of correct patterning during embryogenesis, as they possessed two cotyledons, a hypocotyl, and an identifiable root. As such, they lacked the patterning defects observed as a consequence of a deficient GN function, that is, cotyledon fusion, a compact body pattern without a discernable hypocotyl, and a deletion of roots (Mayer et al., 1991, 1993; Shevell et al., 1994). The absence of single or fused cotyledons is particularly informative, as this phenotype is most sensitive to the interference with GN function, being manifested already in weaker *gn* alleles (Geldner et al., 2004) or in *gn* knockouts partially complemented with GN-GNL1 chimeric ARF-GEFs (Adamowski et al., 2022). Quadruple homozygous *arfa1acfd* seedlings also presented a range of phenotypes, exhibiting a moderate to strong decrease in overall growth rates (Figure 3D). The apical development in these mutants was typically characterized by two cotyledons, which were misshapen, or by a complete apical deletion (Figure 3E). This phenotype contrasts with that resulting from *gn* deficiency, as it classically deletes the basal, rather than the apical, part of the embryonic pattern (Mayer et al., 1991, 1993). Among 725 seedlings of a segregating *arfa1acf* -/-*arfa1d* +/-population, only 5 seedlings with single cotyledons were found, and this phenotype did not correlate with the overall degree of growth deficiency. Genotyping revealed these seedlings to not be *arfa1acfd* quadruple homozygotes (Suppl. Figure 1C). The rarity of this phenotype, and the lack of correlation with *arfa1* genotypes, suggest they represented rare, spontaneous defects in embryonic development, and not the consequences of *arfa1* loss of function. Overall, *arfa1* mutants were not characterized by patterning defects characteristic to *gn* loss of function, which argues against the notion that ARFA1 are targets of regulation by GN through which its developmental patterning function is mediated.

In turn, we identified similarities between *arfa1* mutants and seedlings deficient in GN function in etiolated seedling development. Similarly to *gn* knockouts complemented with partially functional GN-GNL1 chimeras (Adamowski et al., 2022) or to the moderate *GN* mutant allele *gn*^*R5*^ (Suppl. Figure 2A and 2B), both *arfa1acfd* and *arfa1acfe* seedlings grown in the dark variably exhibited agravitropism (Figure 3F), and commonly had open apical hooks after 4 days of culture (Figure 3G). Thus, in contrast with the absence of characteristic patterning defects of embryonic origin, these similarities in post-embryonic growth support a function of ARFA1 in developmental activities mediated by GN.

**Suppl. Figure 2.**
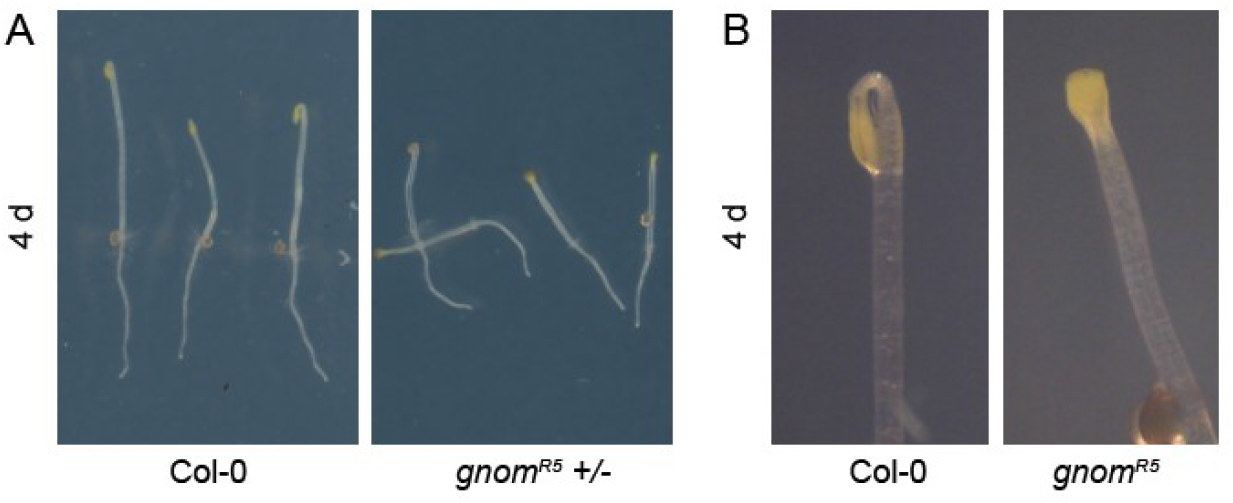
Development of etiolated gnomR5 seedlings. Etiolated gnomR5 seedlings variably exhibit agravitropism (A) and commonly have open apical hooks after 4 days of culture (B).

Overall, the analysis of *arfa1* mutants demonstrates an important role of ARFA1 in general growth and development, but the mutant phenotypes do not clearly indicate their activity as functional partners of GN.

### Cellular phenotypes of *arfa1*

To assess the function of ARFA1 in the endomembrane system, we introduced into *arfa1acf* and *arfa1acf arfa1d* +/- a *δ-COP*_*pro*_*:δ-COP-GFP* reporter construct expressing a subunit of COPI coats, which participate in vesicle formation at the GA (Adamowski et al., unpublished). Using CLSM in seedling RAM epidermis, we observed that individual GA labelled with δ-COP-GFP, normally dispersed in the cytosol, were aggregated into fewer and larger structures in both *arfa1acf* and *arfa1acfd* mutants (Figure 4A). Similar aggregations result from chemical inhibition of ARF-GEFs with Brefeldin A (Robinson et al., 2008), from expression of dominant ARFA1 variants (Singh et al., 2018), and can be observed in mutants of subcellular trafficking components functionally related to the ARF machinery (Zhang et al., 2020). As such, their presence is an indication that endomembrane trafficking is disturbed as a result of ARFA1 deficiency. Expression of δ-COP-GFP also allowed an observation of morphological features of *arfa1* cells and RAM tissues. Some *arfa1acfd* mutant seedlings were characterized by the presence of enlarged vacuoles and very limited volumes of cytosol (Figure 4A, right). Furthermore, *arfa1acfd* mutants with strong phenotypes exhibited deformations of cellular patterns in the RAM, sometimes resulting in hollow spaces in the tissue (Figure 4B). Overall, this live imaging analysis of *arfa1* mutants supports a role of ARFA1 in the endomembrane system activity, previously deduced on the basis of the expression of dominant ARFA1 mutant variants (Lee et al., 2002; Takeuchi et al., 2002; Pimpl et al., 2003; Xu and Scheres, 2005; Singh et al., 2018).

**Figure 4.**
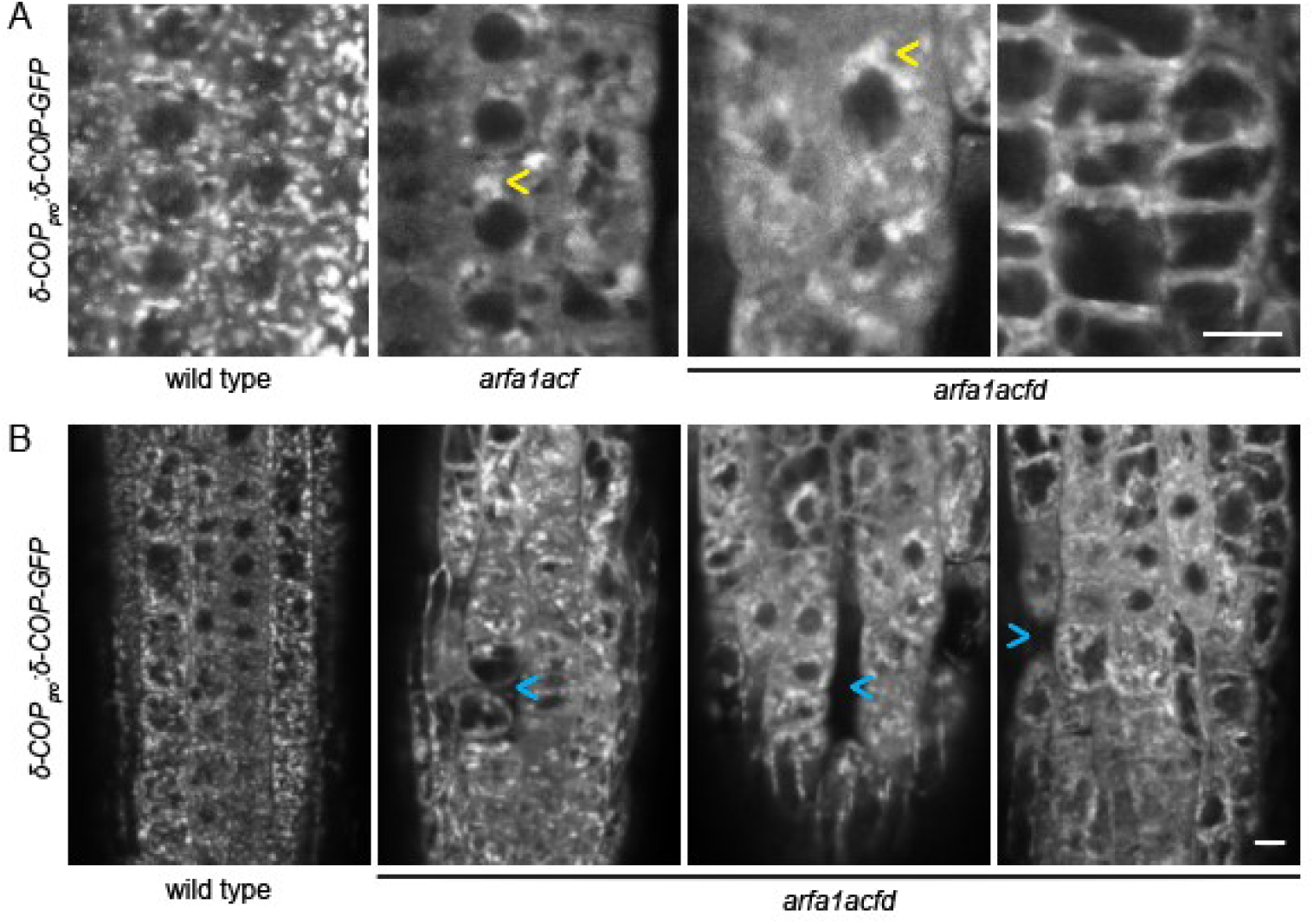
Cellular Phenotypes of arfa1. (A) CLSM images of δ-COP-GFP RAM epidermis of arf1 mutants. GA marked by δ-COP-GFP are aggregated in arf1acf and arf1cafd (arrowheads). Some arf1acdf seedings exhibit an enlargement of vacuoles (rightmost panel). Scale bar - 10 μm. (B) CLSM images of δ-COP-GFP highlighting deformations in cellular patterns in RAM of strong arfa1acfd seedings Arrowheads point to hollow spaces in RAM tissues. Scale bar - 10 μm.

### Subcellular localization of ARFA1 and non-functionality of fluorescent reporters

The *arfa1* mutant phenotypes do not clearly support an ARFA1 function in developmental patterning as targets of GN. The site of action of ARFA1 is established with the use of immunoelectron microscopy (Robinson et al., 2011), which indicates that these ARF isoforms act at the GA and TGN. Yet, functional partners of GN, or of VAN3, are expected to be found at the PM, where these ARF regulators act in their developmental roles (Adamowski et al., 2022; Naramoto and Kyozuka, 2018). To ascertain if the ARFA1 proteins do not localize to the PM, we analysed the previously published *ARFA1c*_*pro*_*:ARFA1c-GFP* line by CLSM in seedling RAMs. Consistent with previous findings, ARFA1c-GFP localized at internal endomembrane system compartments, but not at the PM (Figure 5A). Given that the detection of proteins at the PM of RAM epidermis by CLSM is sometimes limited, we additionally inspected this line by TIRF microscopy in the early elongation zone of the seedling root, but with this method, too, we found no traces of a localization of ARFA1c-GFP at the PM (Figure 5B and Suppl. Movie 2).

**Figure 5.**
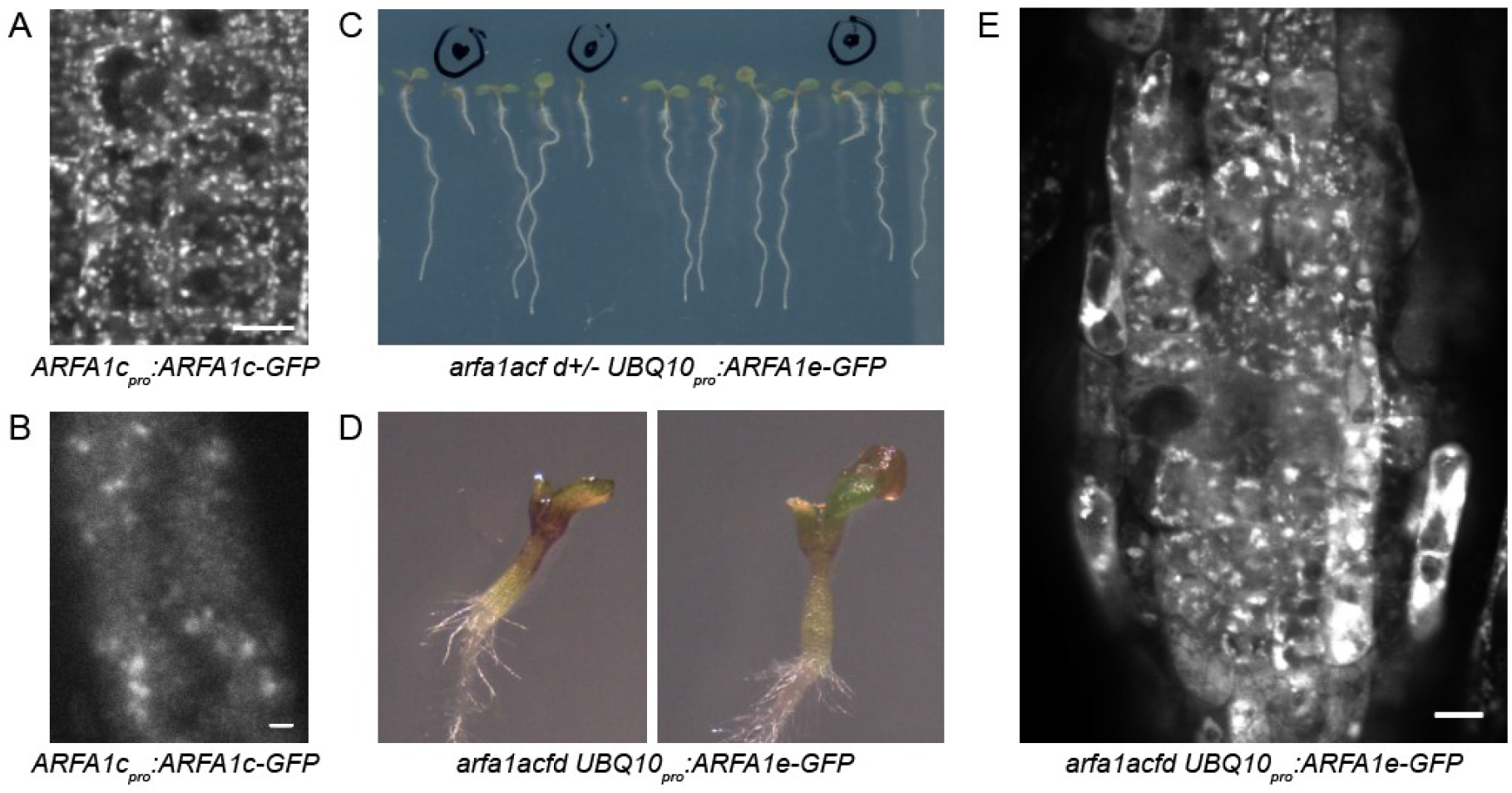
Localization and lack of functionality of ARFA1-GFP fusion proteins. (A) A CLSM image of ARFA1c-GFP in seedling RAM epidermis. Localization to punctate structures, previously identified as GA and TGN, can be observed. No localization to the PM can be observed. Scale bar - 10 μm. (B) A TIRF image of ARFA1c-GFP in the early elongation zone of seedling root. No traces of a localization to the PM is found. Large punctate signals are internal endomembrane structures. Scale bar - 2 μm. See also Suppl. Movie 2 (C) A segregating population of arfa1acf d+/- expressing UBQ10pro:ARFA1e-GFP. Marked seedlings express ARFA1e-GFP and exhibit phenotypes typical for arfa1acfd. (D) Detail of apical parts of arfa1acfd seedlings expressing ARFA1e-GFP. (E) A CLSM image of ARFA1e-GFP expressed in arfa1acfd, demonstrating lack of complementation by the aggregation of endomembrane structures and by the deformities in cellular patterns of the RAM. Scale bar - 10 μm.

To explore this further, we generated a new ARFA1 reporter construct, *UBQ10*_*pro*_*:ARFA1e-GFP*, and introduced it into *arfa1acf* and *arfa1acf arfa1d* +/-lines. Interestingly, this analysis indicated that ARFA1 fluorescent protein fusions are non-functional. Despite ubiquitous expression at high levels, the construct failed to complement the *arfa1* mutant phenotypes at the cellular and morphological levels. In *arfa1acf* background, ARFA1e-GFP localized to endomembrane structures, likely GA and TGN, which appeared to be partially aggregated (Suppl. Figure 1D), as such manifesting the *arfa1acf* cellular phenotype observed with δ-COP-GFP (Figure 4A). In segregating populations of *arfa1acf arfa1d* +/-, seedlings expressing ARFA1e-GFP, but still exhibiting strong *arfa1acfd* phenotypes that were not complemented, were found (Figure 5C and 5D). CLSM imaging of RAMs in these seedlings best demonstrated the lack of ARFA1e-GFP protein fusion functionality: The roots exhibited strongly aggregated compartments labelled by ARFA1e-GFP, and the tissue manifested deformities characteristic for this *arfa1* mutant line (Figure 5E; compare with Figure 4B).

Taken together, the localization of ARFA1 to the GA and TGN is not supportive of their collaboration with GN and VAN3, which, based on current models, act in the developmental patterning function from the PM (Adamowski et al., 2022; Naramoto and Kyozuka, 2018). Yet, fluorescent reporter fusions of ARFA1 cannot be conclusively taken as evidence for a lack of ARFA1 function at the PM, since they are un-functional, and as such, may not necessarily represent all genuine sites of ARFA1 action.

## Discussion

The ARF machinery performs important functions in eukaryotic cells (Jackson and Bouvet, 2014). Much of our information about ARF function in *A. thaliana* comes from the study of ARF-GEF and ARF-GAP regulators, while ARF small GTPases themselves have been studied in comparatively less detail. Of particular interest, the ARF machinery plays an indispensable role in developmental patterning instructed by the hormone auxin, a function associated with the ARF-GEF GN and the ARF-GAP VAN3 (Singh and Jürgens, 2017). We undertook a mutant and localization-based screening of the ARF/ARL family with a primary goal of identifying small GTPases involved in the developmental function carried out by GN and VAN3. The investigation did not yield clear answers with regard to ARF partners that GN and VAN3 might control as mediators of their function. Many of the mutants, even in high order combinations, exhibit no visible phenotypes, arguing against contributions of the mutated genes. Features of the *ARFA1* group, otherwise considered to be the major ARF actor in plants (Singh et al., 2018), are difficult to reconcile with the current concepts about GN and VAN3 function in developmental patterning. GN and VAN3 are both thought to act from the PM (Naramoto and Kyozuka, 2018; Adamowski and Friml, 2021; Adamowski et al., 2022), and while the nature of their activities are unknown, they are expected to involve the control of ARF targets that should be found at the same site of action. Yet, ARFA1 proteins most likely do not localize to the PM, although evaluation through fluorescent protein fusions appears invalid due to their non-functionality. Negative consequences of tagging ARF proteins at the C-terminus are not without precedent (Jian et al., 2010). In any case, ARFA1 localization at the PM was not reported in immunoelectron microscopy studies where native ARFA1 were labelled (Robinson et al., 2011). Beside this inconsistency, the *arfa1* mutants do not manifest a typical *gn* loss of function phenotype of fused or single cotyledons (Mayer et al., 1993; Geldner et al., 2004), but rather possess two cotyledons even when phenotypes are strongly deleterious, or, are characterized by apical deletions with continued development of roots, in direct opposition of *gn* characterized by a lack of root development (Mayer et al., 1991). These are significant arguments against the involvement of ARFA1 in the function of GN, since the loss of ARF partners can be expected to lead to similar consequences as the presumed lack of their GEF-mediated activation. However, similarities between *arfa1* and *gn* were found in the agravitropic growth and deficiencies in apical hook formation in etiolated seedlings, and these appear to support the common involvement of GN and ARFA1 in those aspects of post-embryonic growth. In turn, the severe phenotypes in *arfa1* knockouts demonstrate that ARFA1 are, overall, major ARF players in *A. thaliana*, as previously indicated by other evidences (Gebbie et al., 2005; Singh et al., 2018). They are most likely the main actors in the trafficking functions carried out at the GA and TGN and controlled by the ARF-GEFs GNL1 and BIG and by the ARF-GAPs AGD5-10 (Richter et al., 2007, 2014; Teh and Moore, 2007; Min et al., 2007, 2013; Sauer et al., 2013). It could be perhaps conceived that ARFA1 participate both in these basic trafficking activities, and in the patterning function associated with GN, but as a result of mutation, the fundamental trafficking function becomes deficient to the degree that causes lethality before the consequences of the loss of the patterning function can be clearly manifested. In other words, in this scenario, a putative function of ARFA1 in developmental patterning would require only a very low fraction of the ARFA1 protein pool. Alternatively, it is possible that a larger group of ARF/ARs, including other homologues beside ARFA1, redundantly mediates the function of GN, precluding phenotypic manifestation in *arfa1*. The lack of a PM localization of fluorescent protein fusions of any of the ARF/ARL isoforms except ARFB1a might be – but this is purely a supposition – not necessarily representative of their true site of action, again due to non-functionality. Finally, a scenario can be proposed where the developmental patterning function of GN is not carried out through its molecular activity as an ARF-GEF, and as such, does not involve any ARFs, but instead results from an additional, unknown biochemical activity, designating GN a “moonlighting protein” (Jeffery, 2018). Yet, this proposition, by itself improbable, appears particularly unlikely if VAN3 is additionally taken into consideration. It would be a truly uncanny scenario if two molecularly distinct proteins, both originally ARF regulators, evolved to possess alternative biochemical functions in order to act in a common molecular mechanism not related to ARF small GTPase control.

## Acknowledgements

The authors would like to gratefully acknowledge Ms Rabab Keshkeih for help in cloning ARF sequences, Prof. Ari Pekka Mähönen for sharing the p1R4-pUBQ10:XVE plasmid, and Prof. Qi-Jun Chen for sharing plasmids used for CRISPR/Cas9 mutagenesis.

## Materials and Methods

### Plant material

The following previously described *A. thaliana* lines were used in this study: *arfc1-2* (SALK_027975; Delgadillo et al., 2020), *gnom*_*R5*_ (Geldner et al., 2004), *ARFA1c*_*pro*_*:ARFA1c-GFP* (Xu and Scheres, 2005), *δ-COP*_*pro*_*:δ-COP-GFP* (Adamowski et al., unpublished). The lines generated as part of this study as listed in Suppl. Table 2, primers used for genotyping in Suppl. Table 3, and details of mutant alleles in Suppl. Table 1.

### *In vitro* cultures of *Arabidopsis* seedlings

Seedlings were grown in *in vitro* cultures on half-strength Murashige and Skoog (1/2MS) medium of pH=5.9 supplemented with 1% (w/v) sucrose and 0.8% (w/v) phytoagar at 21°C in 16h light/8h dark cycles with Philips GreenPower LED as light source, using deep red (660nm)/far red (720nm)/blue (455nm) combination, with a photon density of about 140µmol/(m^2^s) +/-20%. Beta-estradiol (Sigma-Aldrich) was solubilized in 100% ethanol to 5 mg/mL stock concentration and added to 1/2MS media during preparation of solid media to a final concentration of 2.5 µg/mL.

### Molecular cloning and the generation of transgenic lines

All constructs were generated using the Gateway method (Invitrogen) and are listed in Suppl. Table 4. DNA sequences were amplified by PCR reactions using iProof High Fidelity polymerase (BioRad). Primers used for cloning are listed in Suppl. Table 3. *ARF* coding sequences without stop codons were amplified and cloned into pDONR221 vectors by BP Clonase. *UBQ10*_*pro*_*:ARF-GFP* and *UBQ10*_*pro*_*:ARF-mCherry* constructs were prepared by combining ARF/pDONR221, UBQ10_pro_/pDONRP4P1r, and GFP/pDONRP2rP3 or mCherry/pDONRP2rP3 plasmids with pB7m34GW destination vector (Karimi et al., 2002) using LR Clonase II Plus. *UBQ10-XVE>>ARFB1a-GFP* and *UBQ10-XVE>>ARFD1a-GFP* were cloned by combining ARF/pDONR221, p1R4-pUBQ10:XVE (Siligato et al., 2016), and GFP/pDONRP2rP3, with pB7m34GW destination vector (Karimi et al., 2002) using LR Clonase II Plus. Col-0 plants were transformed by the standard floral dip method. T1 transformants were selected on solid 1/2 MS medium without sucrose supplemented with Basta.

### CRISPR mutagenesis

CRISPR mutagenesis was performed with the use of pHEE401 binary vector and template plasmids pCBC-DT1T2 and pCBC-DT2T3 (Wang et al., 2015). sgRNA sequences were selected with the use of CRISPR RGEN Tools website (http://www.rgenome.net/cas-designer/). In order to select sgRNAs targeting several homologous *ARF* genes, information about “off-targets” was used to pick sgRNAs with multiple desired targets. All constructs and the sgRNA sequences employed are listed in Suppl. Table 4. In T1 plants, target sequences were PCR-amplified and sequenced using primers listed in Suppl. Table 3. Selected plants homozygous or heterozygous for *arf* mutations were propagated to T2 generation, where individuals negative for the CRISPR/Cas9 transgene were selected based on genotyping with primers CAS9-F and CAS9-R (Suppl. Table 3). The selected individuals were genotyped again for *arf* mutations. In cases where multiple *ARF* genes were targeted, and T1 plants did not carry all intended mutations, CRISPR/Cas9-positive T2 plants were genotyped to find higher order mutants. Due to high number of inserted CRISPR/Cas9 transgenes, *arfa1b* mutants and higher order mutants derived from them by crossing, as well as *arfa1acfe* mutants, carry the CRISPR/Cas9 transgenes.

### Light microscopy

High magnification images of seedlings developing in *in vitro* cultures were taken with Leica EZ4 HD stereomicroscope.

### Confocal Laser Scanning Microscopy

4 to 5 d old seedlings were used for live imaging with Zeiss 800 confocal laser scanning microscope with 20X and 40X lenses.

### Total Internal Reflection Fluorescence microscopy

Early elongation zone of roots in excised ∼1 cm long root tip fragments from 7 d old seedlings, as well as 3 d old etiolated hypocotyls, were used for TIRF imaging. Imaging was performed with Olympus IX83 TIRF microscope, using a 100X TIRF lens with an additional 1.6X magnification lens in the optical path. Time lapses of 100 frames at 1 s intervals with exposure times of 200 ms were taken.

### Accession numbers

Sequence data from this article can be found in the GenBank/EMBL libraries under the following accession numbers: ARFA1a (AT1G23490), ARFA1b (AT5G14670), ARFA1c (AT2G47170), ARFA1d (AT1G70490), ARFA1e (AT3G62290), ARFA1f (AT1G10630), ARFB1a (AT2G15310), ARFB1b (AT5G17060), ARFB1c (AT3G03120), ARFD1a (AT1G02440), ARFD1b (AT1G02430), ARFC1 (AT3G22950), ARF3 (AT2G24765), ARLC1 (AT2G18390), ARLB1 (AT5G52210), ARLA1a (AT5G37680), ARLA1b (AT3G49860), ARLA1c (AT3G49870), ARLA1d (AT5G67560).

## Author Contribution

M.A. and J.F. conceived the study and interpreted results. M.A. and I.M. performed experimentation. M.A. wrote the manuscript.

