## Supplemental Tables for "ARF small GTPases in the developmental function mediated by ARF regulators GNOM and VAN3"

Suppl. Table 1. Nomenclature of *ARF/ARL* genes and details of mutant alleles

| **AGI code** | **Name used in this study** | **Other names** | **Mutant allele details** |
| --- | --- | --- | --- |
| At1g23490 | *ARFA1a* | *-* | 1^st^ exon 1xT insertion  wt …TTTTGTA…  *arfa1a* …TTTTTGTA… |
| At5g14670 | *ARFA1b* | *-* | - |
| At2g47170 | *ARFA1c* | *ARF1, BEX1* | 2^nd^ exon 1xG insertion  wt …TGGGATG…  *arfa1c* …TGGGGATG… |
| At1g70490 | *ARFA1d* | *-* | 1^st^ exon 1xG insertion  wt …CAAGCTC…  *arfa1d* …CAAGGCTC… |
| At3g62290 | *ARFA1e* | *-* | single mutant: 1^st^ exon 1xA insertion  wt …TCAAGCTT…  *arfa1e* …TCAAAGCTT…  high order mutants: 1^st^ exon 1xA deletion  wt …TCAAGCTT…  *arfa1e* …TCAGCTT… |
| At1g10630 | *ARFA1f* | *-* | 1st exon 5 nt deletion  wt …TTTGTACAAGCT…  *arfa1f* …TTTAGCT… |
| At2g15310 | *ARFB1a* | *ARF9, ARFB* | 1^st^ exon 1xA insertion  wt …TGAAGCT…  *arfb1a* …TGAAAGCT… |
| At5g17060 | *ARFB1b* | *ARF7, ARFA* | 3^rd^ exon 17 nt deletion  wt …GGATGTTGGTGGCCAAGAGAAAC…  *arfb1b* …GGAAAC… |
| At3g03120 | *ARFB1c* | *ARF8, ARFA* | 2^nd^ exon 1xA insertion  wt …ACAATCG…  *arfb1c* …ACAAATCG… |
| At3g22950 | *ARFC1* | *ARL5, MTV8, ARFC1a* | 1^st^ exon T-DNA insertion SALK_027975 *(arfc1-2)* |
| At1g02440 | *ARFD1a* | *ARL4* | Two 1xT insertions in 1^st^ and 2^st^ exons  wt …ATTTGCC…  *arfd1a* …ATTTTGCC…  wt …AAATACA…  *arfd1a* …AAATTACA… |
| At1g02430 | *ARFD1b* | *ARL3* | 226 nt deletion spanning exons 1 and 2, flanking sequences:  *arfd1b* …AAAGCCATTT (…) ACAAGGATTC… |
| At2g24765 | *ARF3* | *ARL1, ARFC1b* | 197 nt deletion spanning exons 1 and 2, flanking sequences:  *arf3* …TCTCTTCCGT (…) ATGGGGGAAG… |
| At5g37680 | *ARLA1a* | *ARL8c* | 2^nd^ exon 1xT deletion  wt …TCACTCGT…  *arla1a* …TCACCGT… |
| At3g49860 | *ARLA1b* | *ARL8d* | Large deletion in the locus between 3^rd^ exon of *ARLA1c* and 2^nd^ exon of *ARLA1b,* with an insertion from the pHEE401 T-DNA sequence mapping near LB, containing partial Hygromycin resistance gene and CamV polyA signal. Flanking sequences:  *arla1bc* …GACATGATTC (…) AGACATGAT… |
| At3g49870 | *ARLA1c* | *ARL8a* |  |
| At5g67560 | *ARLA1d* | *ARL8b* | 3^rd^ exon 1xC deletion  wt …agACCGGT…  *arla1d* …agACGGT… |
| At5g52210 | *ARLB1* | *GB1, ARFRP1* | 1^st^ exon T-DNA insertion SALK_127883 *(arlb1)* |
| At2g18390 | *ARLC1* | *ARL2, TITAN5, HALLIMASCH* | - |

Suppl. Table 2. Lines generated as part of this study.

| **Line name** | **Notes** |
| --- | --- |
| *arfa1e* |  |
| *arfa1acf* |  |
| *arfa1acf d*+/- |  |
| *arfa1acfe* | CRISPR/Cas9 positive |
| *arfa1acf* *δ-COP_pro_:δ-COP-GFP* |  |
| *arfa1acf d*+/- *δ-COP_pro_:δ-COP-GFP* |  |
| *arfb1abc* | CRISPR/Cas9 positive |
| *arfd1ab* |  |
| *arf3* |  |
| *arfb1abc arfc1-2* | ARFB1 CRISPR/Cas9 positive |
| *arfb1abc arfd1ab* | ARFB1 CRISPR/Cas9 positive |
| *arfb1abc arf3* | ARFB1 CRISPR/Cas9 positive |
| *arfb1abc arfc1-2 arfd1ab* | ARFB1 CRISPR/Cas9 positive |
| *arfb1abc arfc1-2 arf3* | ARFB1 CRISPR/Cas9 positive |
| *arfb1abc arfd1ab arf3* | ARFB1 CRISPR/Cas9 positive |
| *arfb1abc arfc1-2 arfd1ab arf3* | ARFB1 CRISPR/Cas9 positive |
| *arfc1-2 arf3* |  |
| *arfc1-2 arfd1ab arf3* |  |
| *arla1bc* |  |
| *arla1abcd* |  |
| *arlb1* | SALK_127883 |
| *arfa1acf UBQ10_pro_:ARFA1e-GFP* |  |
| *arfa1acf d*+/- *UBQ10_pro_:ARFA1e-GFP* |  |
| *XVE>>ARFB1a-GFP* |  |
| *UBQ10_pro_:ARFB1b-mCherry* |  |
| *UBQ10_pro_:ARFB1c-mCherry* |  |
| *UBQ10_pro_:ARFC1-GFP* |  |
| *XVE>>ARFD1a-GFP* |  |
| *UBQ10_pro_:ARF3-GFP* |  |
| *UBQ10_pro_:ARLA1a-GFP* |  |
| *UBQ10_pro_:ARLA1b-GFP* |  |
| *UBQ10_pro_:ARLA1c-GFP* |  |
| *UBQ10_pro_:ARLA1d-GFP* |  |
| *UBQ10_pro_:ARLB1-GFP* |  |
| *UBQ10_pro_:ARLC1-mCherry* |  |

Suppl. Table 3. Primers used in this study.

| **Primer name** | **Sequence** | **Purpose** |
| --- | --- | --- |
| ARFA1a-cF | ttcgatttcacctttttctcg |  |
| ARFA1a-cR | atccccaggtagtgatgcaa |  |
| ARFA1c-cF | CGCTTGTTTGTAATGCTGGA |  |
| ARFA1c-cR | CGAAATTCCAGCTGAGTTGA |  |
| ARFA1d-cFn | tggcattatagatgcataatttagga |  |
| ARFA1d-cRn | TCATTCAGCATCCTGTGGAG |  |
| ARFA1e-cF | tgatgcaaaatgagttccaga |  |
| ARFA1e-cR | CAAAAACGAGAAGCACAGCA |  |
| ARFA1f-cF | ttgtcgtcgaatgatcgtgt |  |
| ARFA1f-cR | CAACCGTCTCCACATTAAACC |  |
| ARFB1a-cF | tttccagcctttcttctaatgg |  |
| ARFB1a-cR | cttcctcaccaatgcaaaca |  |
| ARFB1b-cF | atctggatgggatctcgtgt |  |
| ARFB1b-cR | aaagattggccaagcaacat |  |
| ARFB1c-cF | tgatgtagccaatccactttca |  |
| ARFB1c-cR | GAGCATGAATGGGTCCCTTA |  |
| attB1-ARFC1-F | GGGGACAAGTTTGTACAAAAAAGCAGGCTTTATGGGAGCATTCATGTCGA |  |
| arfc1-SALK-R | agcaaacagttcctccggta |  |
| ARFD1a-cF | tgaggcaaaacatcctgtca |  |
| ARFD1a-cR | ggtttctttgcacCTCATCG |  |
| ARFD1b-cF | tgacttttgactttcgcttgc |  |
| ARFD1b-cR | TGTTTCGTCATTGGAAACCA |  |
| ARF3-cF | aatgggccgagctgataata |  |
| ARF3-cR | AAATCCCAGACCTGAAACTTGA |  |
| ARLA1a_F | GGGGACAAGTTTGTACAAAAAAGCAGGCTTTATGGGTCTTTGGGATTCACTT |  |
| ARLA1a-cR | GCCATTTAAAGACGGTTTCG |  |
| ARLA1c-cF | ggcatttggtgtgaccttgt |  |
| ARLA1b-R2 | TTGGAATCAACATGAAACCAAG |  |
| ARLA1d-cF | caatgattggggaaatccaa |  |
| ARLA1d-cR | TCAGGATCAGCAGCATCAAC |  |
| arlb1-1-F | tgcacaatgtgaatgggtct |  |
| arlb1-1-R | aaaggggaaaagccacAGTT |  |
| ARFA1e_F | GGGGACAAGTTTGTACAAAAAAGCAGGCTTTATGGGTCTATCCTTCGGAAA | ARF/ARL cloning |
| ARFA1e_Rns | GGGGACCACTTTGTACAAGAAAGCTGGGTCAGCCTTGTTTGCGATGTTGT |  |
| ARFB1a_F | GGGGACAAGTTTGTACAAAAAAGCAGGCTTTATGGGAGCCAGATTTTCACG |  |
| ARFB1a_Rns | GGGGACCACTTTGTACAAGAAAGCTGGGTCATACCTCGGACCTCGGACC |  |
| ARFB1b_F | GGGGACAAGTTTGTACAAAAAAGCAGGCTTTATGGGTCAAGCTTTTCGTAAG |  |
| ARFB1b_Rns | GGGGACCACTTTGTACAAGAAAGCTGGGTCAAACGAGTGGCCAACCGAT |  |
| ARFB1c_F | GGGGACAAGTTTGTACAAAAAAGCAGGCTTTATGGGTCAAACTTTTCGCAA |  |
| ARFB1c_Rns | GGGGACCACTTTGTACAAGAAAGCTGGGTCAAACGAGGGACCAACTGATG |  |
| ARFC1_F | GGGGACAAGTTTGTACAAAAAAGCAGGCTTTATGGGAGCATTCATGTCGA |  |
| ARFC1_Rns | GGGGACCACTTTGTACAAGAAAGCTGGGTCACTCGTGGCTTTACCGGTAA |  |
| ARFD1a_F | GGGGACAAGTTTGTACAAAAAAGCAGGCTTTATGGGGACGACTCTGGGAA |  |
| ARFD1a_Rns | GGGGACCACTTTGTACAAGAAAGCTGGGTCCATTCTTTCAGCATTTTTCAACA |  |
| ARFD1b_F | GGGGACAAGTTTGTACAAAAAAGCAGGCTTTATGGGGACAGCTCTGGGAA |  |
| ARFD1b_Rns | GGGGACCACTTTGTACAAGAAAGCTGGGTCCATTCTTTCAGCATTTTTCAACA |  |
| ARF3_F | GGGGACAAGTTTGTACAAAAAAGCAGGCTTTATGGGAATCTTATTCACGCG |  |
| ARF3_Rns | GGGGACCACTTTGTACAAGAAAGCTGGGTCGCCACTTCCCGACTTCAAT |  |
| ARLA1a_F | GGGGACAAGTTTGTACAAAAAAGCAGGCTTTATGGGTCTTTGGGATTCACTT |  |
| ARLA1a_Rns | GGGGACCACTTTGTACAAGAAAGCTGGGTCTGTGGCAGTTCTCGAGTGCT |  |
| ARLA1b_F | GGGGACAAGTTTGTACAAAAAAGCAGGCTTTATGACTTTGCAGCCTAATCTTCA |  |
| ARLA1b_Rns | GGGGACCACTTTGTACAAGAAAGCTGGGTCGTTCTTTGATTTTGAGTGATTTACAA |  |
| ARLA1c_F | GGGGACAAGTTTGTACAAAAAAGCAGGCTTTATGGGTTTGTTGGAAGCTTTT |  |
| ARLA1c_Rns | GGGGACCACTTTGTACAAGAAAGCTGGGTCGTTCTTCGACTTTGAATGCTTT |  |
| ARLA1d_F | GGGGACAAGTTTGTACAAAAAAGCAGGCTTTATGGGTTTGTGGGATGCTCT |  |
| ARLA1d_Rns | GGGGACCACTTTGTACAAGAAAGCTGGGTCATTCGATGATTTGGAGTGCTTT |  |
| ARLB1_F | GGGGACAAGTTTGTACAAAAAAGCAGGCTTTATGTTTTCTCTTATGTCTGGACTATGG |  |
| ARLB1_Rns | GGGGACCACTTTGTACAAGAAAGCTGGGTCTGAATTTGGCACAGGAGTGT |  |
| ARLC1_F | GGGGACAAGTTTGTACAAAAAAGCAGGCTTTATGGGACTGTTAAGCATAATCCG |  |
| ARLC1_Rns | GGGGACCACTTTGTACAAGAAAGCTGGGTCGTCAAGCATGTAAATCCTGGAG |  |

Suppl. Table 4. Constructs generated in this study.

| **Plasmid name** | **Notes** |
| --- | --- |
| ARFA1e/pDONR221 | without STOP codon |
| ARFB1a/pDONR221 | without STOP codon |
| ARFB1b/pDONR221 | without STOP codon |
| ARFB1c/pDONR221 | without STOP codon |
| ARFC1/pDONR221 | without STOP codon |
| ARFD1a/pDONR221 | without STOP codon |
| ARF3/pDONR221 | without STOP codon |
| ARLA1a/pDONR221 | without STOP codon |
| ARLA1b/pDONR221 | without STOP codon |
| ARLA1c/pDONR221 | without STOP codon |
| ARLA1d/pDONR221 | without STOP codon |
| ARLB1/pDONR221 | without STOP codon |
| ARLC1/pDONR221 | without STOP codon |
| UBQ10:ARFA1e-GFP/pH7m34GW |  |
| UBQ10-XVE>>ARFB1a-GFP/pB7m34GW |  |
| UBQ10:ARFB1b-mCherry/pB7m34GW |  |
| UBQ10:ARFB1c-mCherry/pB7m34GW |  |
| UBQ10:ARFC1-GFP/pB7m34GW |  |
| UBQ10-XVE>>ARFD1a-GFP/pB7m34GW |  |
| UBQ10:ARF3-GFP/pB7m34GW |  |
| UBQ10:ARLA1a-GFP/pB7m34GW |  |
| UBQ10:ARLA1b-GFP/pB7m34GW |  |
| UBQ10:ARLA1c-GFP/pB7m34GW |  |
| UBQ10:ARLA1d-GFP/pB7m34GW |  |
| UBQ10:ARLB1-GFP/pB7m34GW |  |
| UBQ10:ARLC1-Cherry/pB7m34GW |  |
| ARFA1 CRISPR 1/pHEE401 | 1^st^ round of mutagenesis  1 AGCTTGAGCTTGTACAAAA  2 TCTTGTACAAGCTCAAGCT  3 TGACCCCCAACATCCCACA |
| ARFA1 CRISPR 3/pHEE401 | 2^nd^ round of mutagenesis  1 TCTTGTACAAGCTCAAGCT  2 TCTTTACCAATAGTAGGGA  3 TGACTACGATTCCAACCAT |
| ARFB1 CRISPR/pHEE401 | 1 TCCTTTACAAGTTGAAGCT  2 TGTCTACTGTTCCCACAAT  3 TGTTCACAGTTTGGGATGT |
| ARFD1 CRISPR/pHEE401 | 1 CTCTGGGAAAGCCATTTGC  2 GTGGAAAGTGTAAAATACA |
| ARF3 CRISPR/pHEE401 | 1 GGATGTTCTCTTCCGTCTT  2 TCCAGATCGGCTTCAGATG |
| ARLA1/ARL8 CRISPR 1/pHEE401 | 1^st^ round of mutagenesis  1 CATTTCTTGCTTGAAGAAG  2 AGTGAAGACATGATTCCTA  3 ACATGAGGAAAGTTACAAA |
| ARLA1/ARL8 CRISPR 2/pHEE401 | 2^nd^ round of mutagenesis  1 AGATGGAGCTGTCACTCGT  2 TTCACTGTATCCACCGGTC |
